# Role of extracellular mycobacteria in blood-retinal barrier invasion in a zebrafish model of ocular TB

**DOI:** 10.1101/2021.01.08.425865

**Authors:** Santhosh Kumar Damera, Ranjan K. Panigrahi, Sanchita Mitra, Soumyava Basu

**Author notes:** **Corresponding author**: Soumyava Basu, L V Prasad Eye Institute (MTC Campus), Patia, Bhubaneswar, India, Email.

## Abstract

Intraocular inflammation following mycobacterial dissemination to the eye is common in tuberculosis (TB)-endemic countries. However, the early host-pathogen interactions during ocular dissemination are unknown. In this study, we investigate the early events during mycobacterial invasion of the blood-retinal barriers (BRB) with fluorescent tagged *Mycobacterium marinum* (*Mm*), host macrophages and retinal vasculature in a zebrafish model of ocular TB. We found that *Mm* invaded the vascular endothelium either in extracellular or intracellular (inside phagocytes) state, typically 3-4 days post-injection (dpi). Extracellular *Mm* are phagocytosed in the retinal tissue, and progress to form a compact granuloma around 6 dpi. Intracellular *Mm* crossing the BRB are likely to be less virulent, and either persist inside solitary macrophages (most cases), or progress to loosely arranged granuloma (rare). The early interactions between mycobacteria and host immune cells can thus determine the course of disease during mycobacterial dissemination to the eye.

## Introduction

Ocular tuberculosis (TB) is a sight-threating form of intraocular inflammation that is common not only in TB-endemic countries but is also being increasingly reported from non-endemic countries [1–2]. Despite its common occurrence, the pathogenesis of ocular TB has remained unclear [3]. Most of our understanding of the pathogenesis of this condition has emerged from histopathological studies of end-stage disease that required enucleation of the eyeball [4–5]. These studies have revealed the presence of granulomatous inflammation and acid-fast bacilli within ocular tissues, both of which unequivocally point towards ocular dissemination of *Mycobacterium tuberculosis* (*Mtb*) in this disease. However, these studies do not provide any information on the early events during mycobacterial dissemination to the eye. The early events are critical as they have the potential to either eliminate the infection completely or limit the establishment and/or expansion of granuloma, thereby influencing the clinical manifestation of the disease [6].

The zebrafish larva provides an opportunity to visualise the early host-pathogen interactions through fluorescent tagged mycobacteria and host-immune cells [7–8]. It has several unique advantages, the most important being its transparency which makes it amenable to live microscopy. The zebrafish and its natural pathogen, *Mycobacterium marinum* (*Mm*), have provided remarkable insights into possible mechanisms of host immune response and granuloma formation during human mycobacterial infection [9–10]. The zebrafish model has also been used to investigate mycobacterial infection in the central nervous system (CNS), especially traversal of mycobacteria across the blood-brain barrier (BBB) [11–13]. It has also been used to study host pathogen interactions in other CNS infections such as *Cryptococcus neoformans* [14].

Recently, we reported a zebrafish model of ocular TB, in which we demonstrated mycobacterial localisation and granuloma formation in the vicinity of blood-retinal barriers (BRB), following injection into the caudal veins of embryos [15]. We also demonstrated recruitment of peripheral blood monocytes into these granuloma suggesting breakdown of the BRB by the inflammatory focus. However, we did not investigate the events surrounding the mycobacterial traversal across the BRB. In this study, we have further explored the mechanisms of mycobacterial invasion of the BRBs and the influence of bacterial virulence on these events.

## Results

### *Mm* causes high rate of ocular infection even with low inoculum of systemic infection

We obtained an ocular infection rate of 60% (18 of 30 infected larvae, in two separate experiemtns) at 1 dpi (figure 1A). The rate of ocular infection was defined as the percentage of *Mm*-injected fish showing presence of *Mm* in at least one of the eyes at 10x magnification under the fluorescent microscope. This remained between 60-64% till 6 dpi, even though there was a steady decrease in viable embryos due to the *Mm* infection. Notably, we used 25 CFU inoculum for caudal vein injections and injected the larvae at 4 dpf, since the inner and outer BRBs as well as the BBB, have been found to established at 3 dpf in earlier studies [16,17]. The 4 dpf time point was also used for caudal vein infection in the TB meningitis model [11,12]. However, the inoculum used in earlier models of ocular TB and TB meningitis ranged between 100-1000 CFU of *Mm* [12,15]. The lower inoculum used in our study can be expected to mimic human extra-pulmonary dissemination better than previous studies.

**Figure 1:**
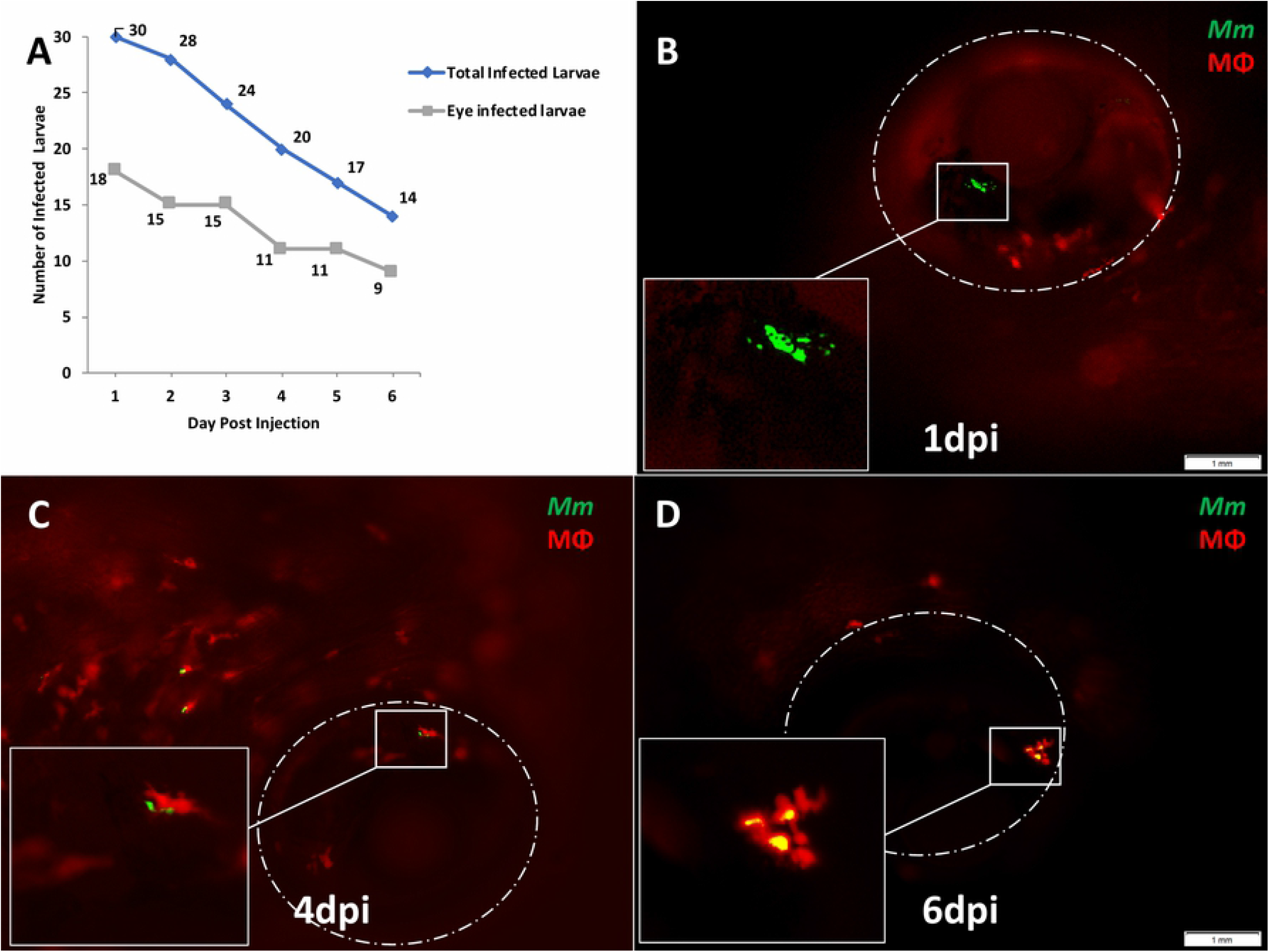
Ocular infection and granuloma formation with wild-type *Mycobacterium marinum* (*Mm*) in *mpeg1:BB* transgenic larvae. (A) Frequency of ocular infection following caudal vein infection with 25 colony-forming units of *Mm*, between 1-6 days post injection. **Granuloma formation was seen in 6 (33.3%) of the 18 eye-infected embryos.** The kinetics of granuloma formation in all the six eyes were as follows: (B) Extracellular *Mm* (green) seeded in the eye at 1 dpi. The *Mm* remained extracellular till 3 dpi. (C) *Mm* phagocytosed inside a solitary macrophage (red) at 4 dpi. (D) Compact granuloma formation at 6 dpi, comprising of macrophages with or without phagocytosed *Mm*.

Next, we evaluated *Tg(mpeg1::BB)* larvae with WT *Mm* ocular infection, for the kinetics of granuloma formation in the eye. Our earlier study had demonstrated granuloma comprising of aggregates of infected macrophages inside ocular tissues of infected zebrafish larvae [15]. The macrophages in these granulomas appeared to be derived from the circulating monocytes as well as from the resident macrophage population within the retina. In the current study, granuloma formation was seen in 6 (33.3%) of the 18 eye-infected embryos. In ocular infections that progressed to granuloma formation, we noted that between 1 and 3 dpi, the *Mm* remained extracellular either as single or multiple clumps (figure 1B). No phagocytosis was noted till 3 dpi. In these larvae, phagocytosis of the ocular *Mm* was first noted at 4 dpi in four eyes and at 5 dpi in the remaining two (figure 1C). Additional macrophages were seen in the vicinity of infected macrophages on 5 dpi, probably as a result of chemotaxis (data not shown). Finally, on 6 dpi, aggregation of macrophages into granulomas was noted (figure 1D). The granulomas were typically compact with tight arrangement of macrophages, with or without *Mm* inside them. A different temporal sequence was noted in four of the remaining 12 embryos. Here, phagocytosis was first seen at 3 dpi, and the *Mm* continued to remain inside solitary macrophages without progression to granuloma formation till 6 dpi (figure 2A-D). In the remaining eight larvae, the *Mm* were not seen beyond 3 dpi in the eyes. These Mm could have moved out of the ocular circulation (as noted in our earlier study), or may have been removed by circulating phagocytes.

**Figure 2:**
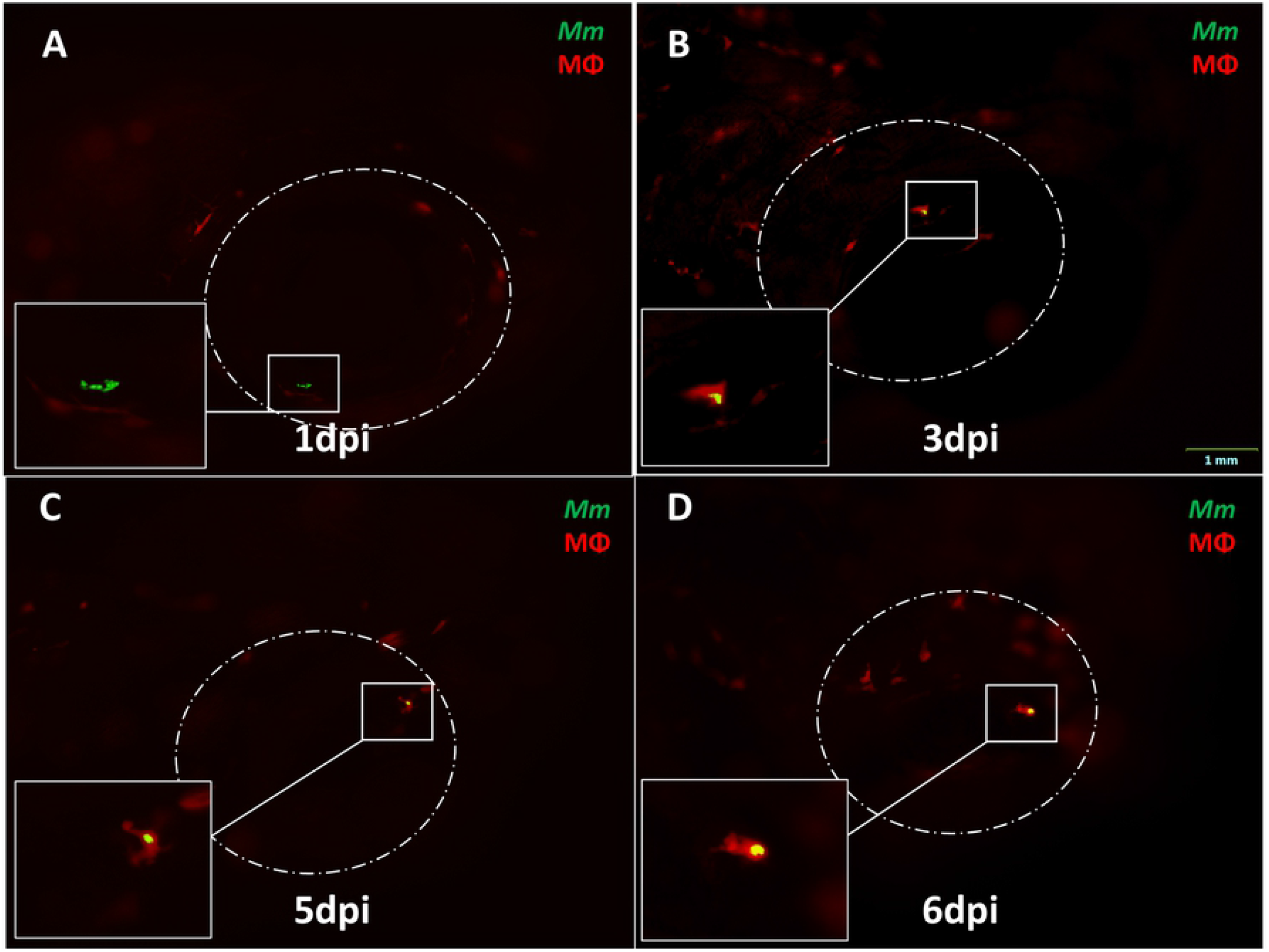
Progression of ocular infection in eyes with early phagocytosis in *mpeg1:BB* transgenic larvae. (A) Extracellular *Mycobacterium marinum* (*Mm*, green) in the eye at 1 dpi. (B) Phagocytosis (red macrophages) is seen early at 3 dpi. (C-D) *Mm* remain inside solitary macrophages till 6 dpi, with no evidence of aggregation into granuloma.

The dichotomy noted in the course of events between extracellular *Mm* versus those that were phagocytosed early at 3 dpi, led us to suspect that phagocytosis influenced the traversal across the BRB and progression to granuloma formation. To further dissect the kinetics of infection at the BRB (retinal vascular endothelium), we infected double transgenic (cross bred for *kdrl* and *mpeg1: BB*) larvae with WT *Mm*. We noted that till 3 dpi, 83.3% (10 of 12) of the *Mm* were extracellular and within the lumen of the blood vessel (figure 3A-B). We also found *Mm* in the wall of the vessel at 3 dpi (figure 3B). Only two *Mm* (both of which were extracellular) were also found to have already crossed over to the retinal tissue (non-vascular region). The first evidence of phagocytosis was noted at 4 dpi only after entering the retinal tissue, and only solitary macrophages with engulfed *Mm* were seen till 5 dpi (figure 3C). Aggregation of macrophages into nascent granuloma was seen only at 6 dpi (figure 3D).

**Figure 3:**
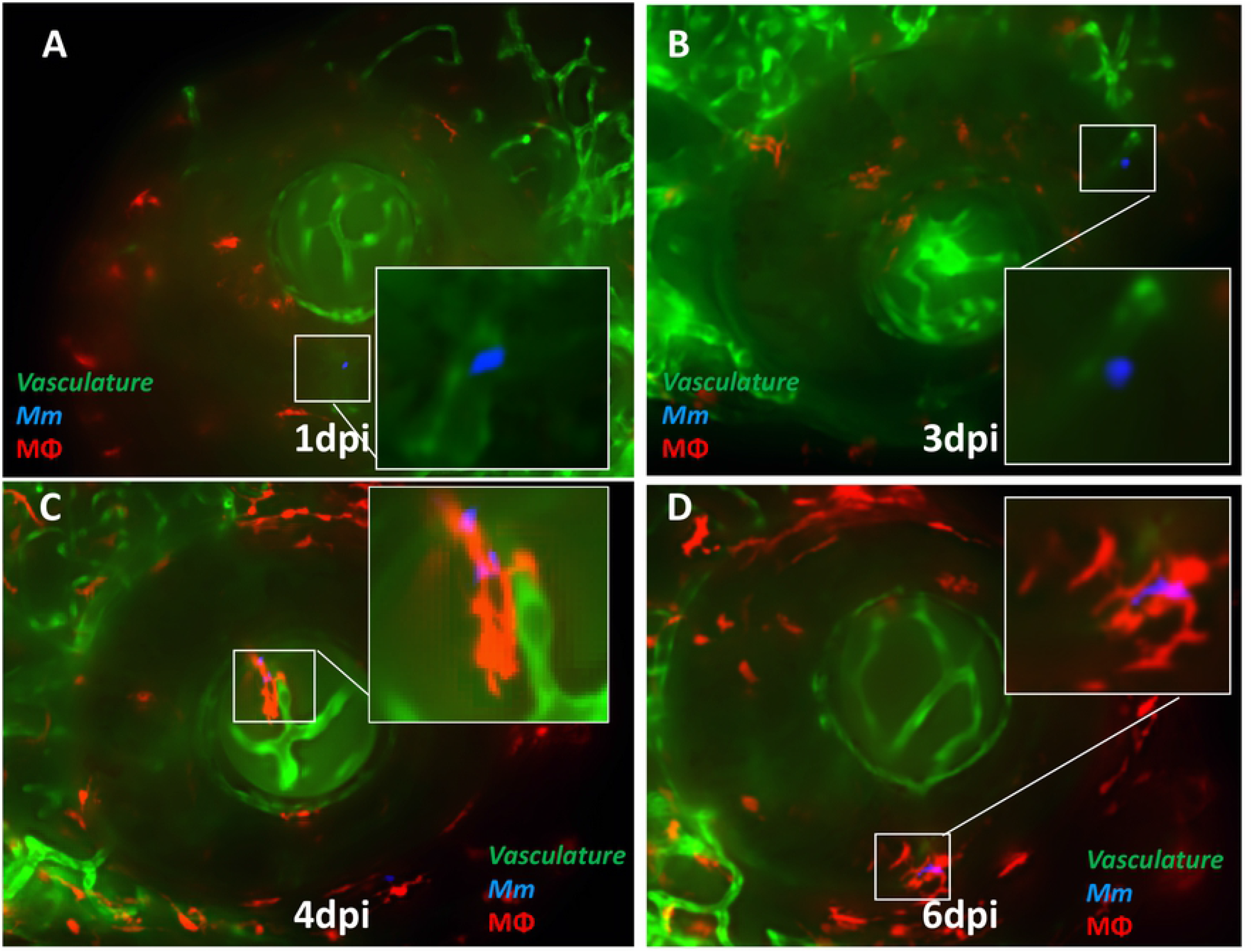
Kinetics of ocular infection and granuloma formation in double transgenic (cross bred for *kdrl* and *mpeg1: BB*) larvae with wild-type *Mycobacterium marinum* (*Mm*). (A) Extracellular *Mm* (blue) lying in the lumen of retinal blood vessels at 1 dpi, and (B) crossing the vascular endothelium at 3 dpi. (C) The first appearance of phagocytosis inside the retinal tissue at 4 dpi, with the infected macrophage closely abutting the vascular endothelium, and (D) Aggregation of macrophages into a granuloma within the retinal tissue at 6 dpi.

### Depletion of circulating monocytes increases rate of ocular infection

Intracellular organisms such as mycobacteria or cryptococci are known to rely primarily on professional phagocytic cells such as macrophages and dendritic cells for crossing vascular endothelial barriers such as the BBB or the BRB [12,14,18]. This is known as the Trojan Horse mechanism and has been elegantly described in zebrafish models of central nervous system infection such as TB, and cryptococcus [12,14]. We therefore hypothesised that depletion of the circulating macrophages with liposomal clodronate will reduce the rate of ocular infection especially beyond 3 dpi, when the crossing across the BRB typically happens. To confirm the efficacy of the drug we found that the number of circulating macrophages was significantly lower on all days till 6 dpi, compared to larvae treated with control liposomes (figure 4A-C). Surprisingly, the rate of ocular infection in clodronate treated larvae remained consistently higher than controls, even beyond 3 dpi, when the *Mm* crosses the BRB (figure 4D). The mean percentage of larvae with eye infection in the clodronate treated group (49.98±9.57) was significantly more than the control group (38.73±8.46) over 6-days post injection (p<0.03, student t-test). However, the rate of granuloma formation amongst the infected larvae was marginally lower in the clodronate treated group (8%, 2 of 25 larvae) as compared to the control group (14.3%, 4 of 28 larvae). We expect that the lack of contribution from circulating macrophages could be responsible for the lower rate of granuloma formation.

**Figure 4:**
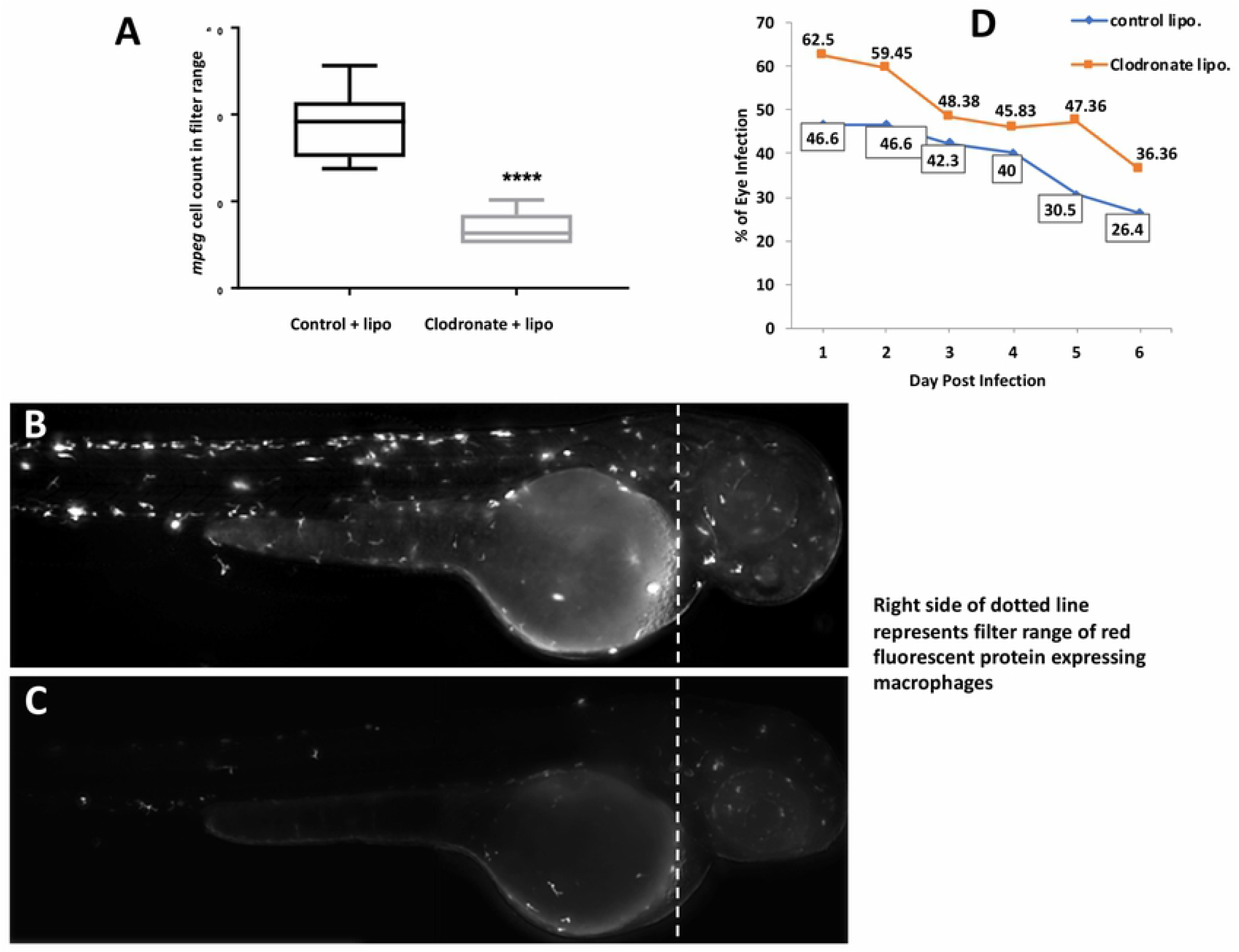
Effect of depletion of circulating monocytes on ocular infection. Liposomal clodronate was injected into *mpeg1:BB* transgenic larvae at 3 days post fertilization (DPF), followed by wild type *Mycobacterium marinum* injection at 4 dpf. Monocyte depletion within the head portion (shown by dashed line in B and C) was confirmed by loss of red fluorescence within the filter range at 4 dpf (Cellsense mean fluorescence intensity [MFI}) and additionally by counting manually under fluorescent microscopy. (A) MFI within the selected head area was significantly lower in the clodronate liposome treated larvae, as compared to controls on days 1-6 post injection (dpi). **** *p*<0.0001 (B) Representative fluorescent image (2x magnification) of control, and (C) clodronate liposome treated larvae (D) Higher ocular infection rate in clodronate liposome treated larvae as compared to controls on all days from 1-6 dpi. The mean percentage of larvae with eye infection in the clodronate treated group (49.98±9.57) was significantly more than the control group (38.73±8.46) over 6-days post injection (p<0.03, student t-test).

### ESX-1 secretion system is required for crossing BRB and granuloma formation

Next, we sought to determine the influence of bacterial virulence factors in crossing the BRB and subsequent granuloma formation. Here, we used RD1 mutant *Mm* [8,19] that lacks the ESX-1 gene locus encoded secretory system required for secretion of several proteins including Early Secreted Antigenic Target-6 (ESAT-6). Previous studies have demonstrated the role of ESX-1 in invasion of brain endothelial cells during macrophage independent crossing of the blood-brain barrier [12, 19]. Expectedly, the frequency of ocular infection by RD1 mutant *Mm* remained low, ranging between 16.6% to 9.1% over a follow up period of 10 days post-injection (figure 5A). Larvae infected with mutant *Mm* survived longer (up to 15 dpi), likely due to the lower virulence of the infecting organism. We also repeated the macrophage depletion experiment with RD1 mutant *Mm*. Here, we noted that in contrast to WT *Mm*, the rate of infection was higher in the control group as compared to the macrophage depleted group, from 1 till 6 dpi (figure 5B). This observation suggests that macrophages are vital for BRB crossing of less virulent *Mm*.

**Figure 5:**
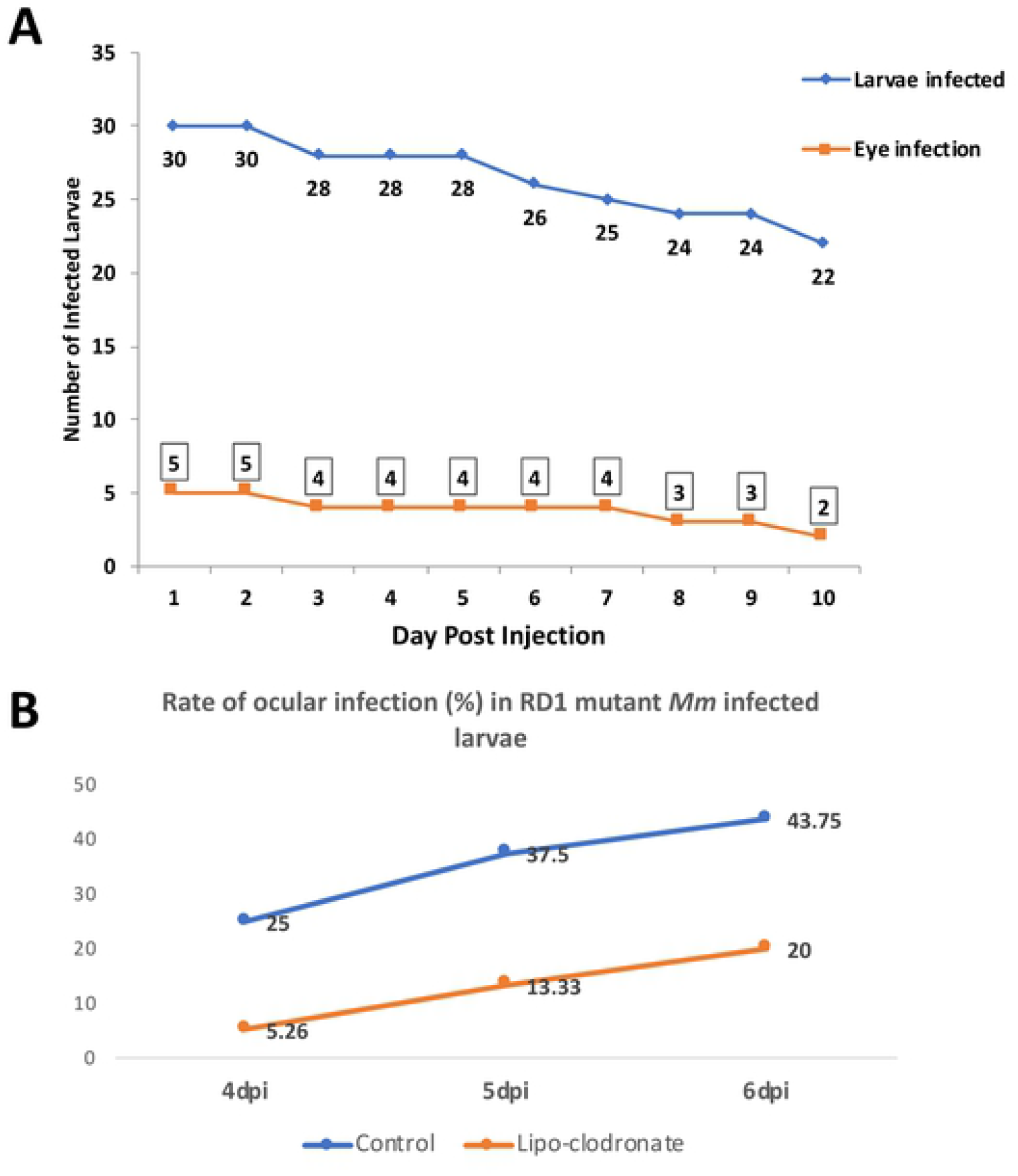
Effect of RD1 mutation in *Mycobacterium marinum* (*Mm*) on frequency on ocular infection. (A) Line graph showing significantly low frequency of ocular infection with RD1 mutant *Mm* (compare with figure 1A), even up to 10 dpi. The infected larvae however survived longer than during infection with wild-type (WT) *Mm*, as shown by the low slope of the graph (B) Depletion of circulating monocytes with clodronate liposomes led to drop in ocular infection rate, the effect being opposite to infection with WT *Mm*.

Finally, we investigated if the absence of ESX-1 secretion system influenced the kinetics of BRB traversal and granuloma formation in double transgenic larvae. We noted a delay among mutant *Mm* in the crossing of BRB (4-5 dpi) as well as in granuloma formation (7 dpi). In fact, only one of the 5 ocular infections showed some signs of aggregation though it need not necessarily be described as a granuloma. The macrophages within the granuloma appeared to be loosely arranged unlike the compact arrangement noted with WT *Mm* (figure 6A-D). To further establish this differential transit across the BRB, we injected equally mixed population of WT and RD1 mutant *Mm* in *kdrl* larvae, where we again found that at 4 dpi, WT *Mm* had already crossed the BRB in 52.9% (9/17) of ocular infections, while RD1 mutant *Mm* continued to be in the blood vessel in all larvae with ocular infections (n=4) (Fig 7).

**Figure 6:**
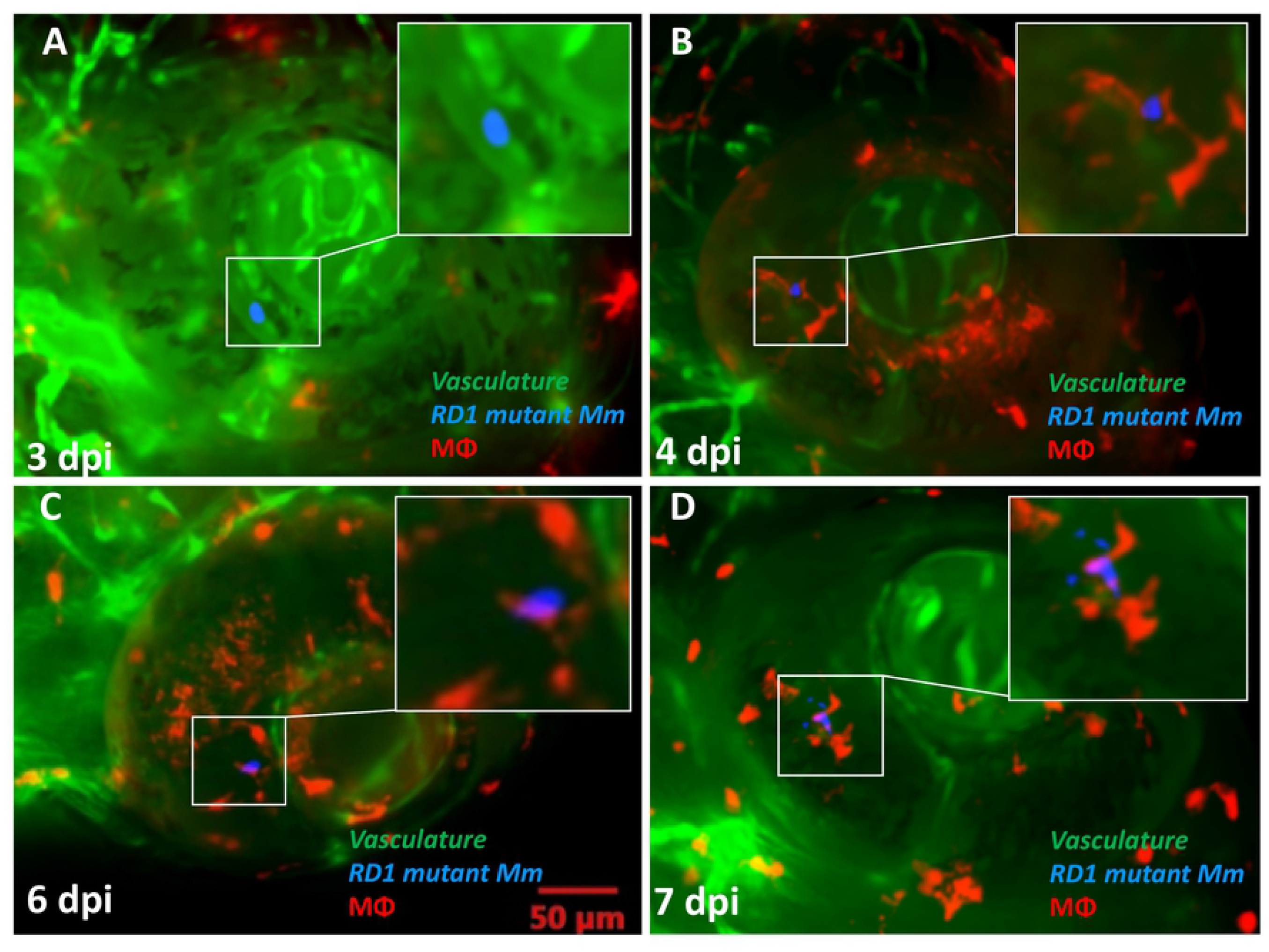
Kinetics of granuloma formation following ocular infection with RD1 mutant *Mycobacterium marinum* (*Mm*). (A) Extracellular Mm (blue) seen within retinal vascular lumen at 2 dpi. (B) Phagocytosed Mm (red macrophage) within the retinal tissue at 4 dpi. (C) Persistence of solitary macrophage with no evidence of aggregation even at 6 dpi. (D) Loose macrophage aggregation at 7 dpi, not representative of a true granuloma. This was seen in only one of the five ocular infections with RD1 mutant *Mm*.

**Figure 7:**
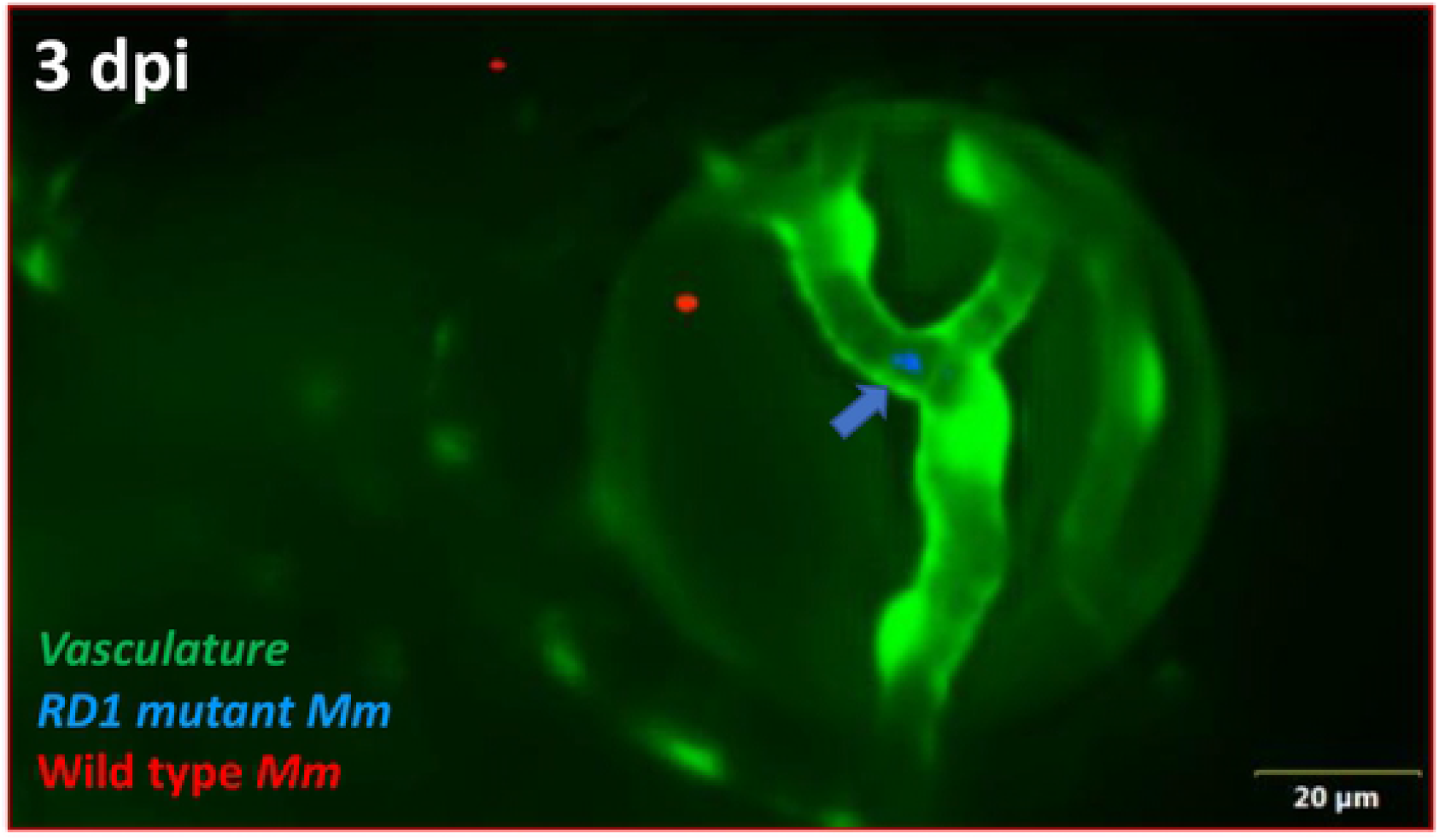
Differential progression of ocular infection by wild type (WT) and RD1 mutant *Mycobacterium marinum* (*Mm*). At 3 dpi, WT *Mm* (red) have already crossed the vascular endothelial barrier and reached the retinal tissue, while RD1 mutant *Mm* (blue, arrow) continue to be inside the vascular lumen (n=4/17).

Collectively, our data suggests an apparent dichotomy in the predominant mechanism of BRB transfer between virulent and avirulent bacteria. Virulent (RD1-competent) *Mm* appear to cross the BRB as free, extracellular bacteria by ESX-1 dependent mechanisms and progress to formation of compact granuloma. Avirulent or less virulent *Mm* require prior phagocytosis by circulating macrophages and cross the BRB by the Trojan Horse mechanism. These infections mostly do not progress to granuloma formation, and even if they do, typically result in loosely arranged granuloma.

## Discussion

CNS invasion is generally caused by intracellular pathogens. Since the Trojan Horse hypothesis adequately explains their ability to cross the CNS barriers, the extracellular pathways of these pathogens have mostly been ignored. This despite the fact that the same pathogens while free or extracellular can cross the CNS barriers nearly as much, or sometimes more efficiently, than while inside phagocytes [12]. In this study, while examining the interactions between mycobacteria and the BRB in a zebrafish model of ocular TB, we demonstrated that virulent (RD1-competent) mycobacteria typically adopt extracellular pathways and progress to granuloma formation in the eye. Avirulent bacteria require the Trojan Horse to get past the BRB, and generally do not progress to granuloma formation.

The role of extracellular Mtb in CNS or retinal invasion is supported by several factors. Recent *in vivo, in vitro* and transcriptional studies suggest the possibility of extracellular dissemination of *Mtb* from the lungs [20–22]. *Mtb* is also able to survive and replicate *ex vivo* in blood where it develops enhanced virulence [23]. *In vivo* and *in vitro* studies have demonstrated that free *Mtb* invade vascular endothelial cells aided by secretion of virulence factors, such as ESAT-6 [12, 19]. Mtb is also able to form a replicative niche within lymphatic endothelial cells [24]. Finally, the likelihood of extracellular dissemination of *Mtb* within the host can also be explained from the evolutionary perspective. The life cycle of *Mtb* depends on causing pulmonary disease and dissemination to a new host [25]. Therefore, it is unlikely that *Mtb* will adopt survival strategies within the phagocyte, for the purpose of extrapulmonary dissemination.

Our data supporting hematogenous dissemination by extracellular *Mtb*, is congruous with earlier studies on the zebrafish embryo model. For example, in the TB meningitis model, *Mm* were found in the brain tissue in all embryos, when phagocytes were similarly depleted [12]. In contrast, phagocyte depletion with the *pu. 1* morpholino resulted in decreased ‘extravascular’ migration of *Mm* in another study [26]. However, in this study, the BRB/BBB that possess tight junctions were not specifically evaluated and therefore, the results are not truly divergent from our study. Our data is also not in conflict with the possibility of infected macrophages from the primary granuloma facilitating its expansion by recruiting new macrophages and also disseminating to deeper tissues [27], or even to distal locations [28]. Either process does not necessarily involve migration across the BRB/BBB which is the primary context of our study. Other than BRB migration, we found that ESX-1 also influences the likelihood of granuloma formation within the retina. Here again, our results conform with earlier reports of virulent mycobacteria using ESX-1 for recruitment of new macrophages to the nascent granuloma, as well facilitating their infection and mobility within the granuloma, thus facilitating their expansion [27].

Our results would have implications on the early events that decide the clinical phenotype and the time to onset of disease in ocular TB. Ocular TB has distinct phenotypes that are known to affect tissues that are either inside or outside the BRB though occasionally both could be affected [29]. The former includes retinitis and retinal vasculitis while choroiditis, cyclitis and iritis are examples of the latter. Based on our results, we would expect that virulent, extracellular bacilli would cross the BRB more often while the less virulent or less invasive bacterial phenotypes would either remain outside the BRB or cross the BRB packaged inside macrophages. Alternatively, relatively avirulent phenotypes in a given host, may enter non-professional phagocytes such as the retinal pigment epithelium (RPE), where they may remain ‘silent’ for a prolonged period. The possibility of ‘silent’ infection, at least in certain phenotypes of ocular TB, is supported by recent *in vitro* studies in the RPE cells. *Mtb* infection in RPE cells *in vitro* revealed that bacterial virulence genes such as ESX-1 are down-regulated within infected RPE cells [30]. RPE cells not only limit intracellular *Mtb* replication, but also survive longer in the presence of infection [31]. We speculate that virulent transformation of such avirulent phenotypes on a later date, could lead to initiation of the inflammatory cascade in such eyes.

Our interpretation of the results of this study are limited by the absence of two important links. First, we could not colocalise phagocytosed (intracellular) *Mm* within the GFP-expressing endothelial cells during the stage of BRB invasion thus missing out on the actual event of crossing by extracellular *Mm*. Second, we could not demonstrate fluorescent dye (FITC-dextran/ sodium fluorescein) leakage at the point of contact of *Mm* with the vascular endothelium. The larval coats were so thick at 3 dpi (7 dpf) that they did not allow entry of the injecting needle into the caudal vein.

Nevertheless, our study provides reasonable evidence of the likelihood of dichotomous pathway with distinct host responses, during mycobacterial invasion of the BRB. Our observations should also provide direction to the investigation of CNS barriers, in meningeal TB as well as in other infections of the CNS.

## Methods

### Zebra Fish Husbandry

Wild type AB and various transgenic (Tg) zebrafish *Tg(kdrl::EGFP*, green fluorescent endothelial cells), *Tg (mpeg1::BB*, red fluorescent macrophages) and double *Tg (KDRL::mpeg1*, red fluorescent macrophages with green fluorescent endothelial cells) were maintained as previously described [8,32]. Eggs were obtained by natural spawning and fertilized embryos were treated with 0.2mM 1-phenyl-2-thiourea (PTU) (Sigma-Aldrich, P7629) to prevent melanophore formation. The guidelines recommended by the Committee for the Purpose of Control and Supervision of Experiments on Animals (CPCSEA), Government of India, were followed for zebrafish maintenance and experimentation.

### Bacterial Strains

Wild type *Mm* (WT/pmsp12: wasabi), WT/ pmsp12::tdTomato (pTEC27 deposited with Adgene), WT/pmsp12::EBFP2, expressing green, red and blue fluorescent proteins respectively, and RD1 mutant described by Volkman et al. [8,32], were grown at 32° C in Middlebrook’s 7H9 broth (M198, Himedia, Mumbai, India) supplemented with Middlebrook oleic acid/albumin/dextrose/catalase (OADC) and antibiotic hygromycin B (final concentration 50 mg/mL). Culture and preparation of single cell *Mm* for infections were made as described [8], and were stored at final dilution 25 colony forming units (CFU)/nL in 5μL and 10μL aliquots respectively, at −80° C.

### Caudal vein Injections

Dechorionated larvae were anesthetized with 0.02% (w/v) ethyl 3-aminobenzoate methanesulfonate (MS-222) (Tricaine, Sigma-Aldrich, A5040) supplemented with 0.2mM PTU. A single aliquot of bacterial suspension was thawed and mixed with 0.25μL of 20% phenol red as described [8]. For every new aliquot, the bacterial count per nL, was checked under the fluorescence microscope and the colony count checked by plating on Middlebrook’s 7H10 medium. It was also ensured that the same size of drop is used in all injections without changing the injector settings. At 4 days-post-fertilization (dpf), larvae were infected via caudal vein by injecting 1-2nL single cell suspension containing ~25 preprocessed *Mm*. Bacterial injections were controlled using an Olympus SZ stereo-microscope with a Femtojet micro-injector (Eppendorf). Injected larvae were recovered by transferring them into fresh embryo media supplemented with 0.2mM PTU. Physically damaged/ non-viable embryos were discarded prior to injections. During the course of follow up, larvae were transferred into fresh embryo media containing PTU, every 24 hours.

### Microscopy

For live imaging, larvae were anesthetized with 0.02% (w/v) tricaine and screened for eye infection using Olympus BX53 upright fluorescent microscope with z-stack attachment and Olympus Cellsense imaging software. For the infection analysis, larvae were anesthetized and mounted in 3% methyl cellulose and manipulated to place them laterally on flat side to view one eye at a time [15]. Imaging was done with 20x (U PlanFL N, 20x/0.50) objective along with FITC/TRITC/DAPI filters to detect green/red/blue fluorescent light respectively. Presence of bacteria and infected eyes were scored manually, and early infection and granuloma was confirmed and analyzed visually. The primary outcome measures were frequency of ocular infection, timing of phagocytosis and aggregation of macrophages into a granuloma.

### Depletion of circulating monocytes

To deplete circulating monocytes selectively, 2-3nL clodronate liposomes and phosphate buffered saline (PBS) loaded control liposomes with 0.1% phenol red suspension was injected intravenously in the caudal vein of 3 dpf *Tg* (*mpeg1: BB*) larvae [33]. We did not use any anti-sense oligonucleotide (morpholino) directed against the myeloid transcription factor gene pu.1, to prevent overwhelming infection and early death of the embryos. Additionally, persistence of tissue resident macrophages in the retina, allowed us to visualize the post-BRB migration events up to 6 dpi. At 1 day-post-injection (dpi) monocyte depletion in clodronate liposomes injected larvae were confirmed by loss of red fluorescence within the filter range (Cellsense mean fluorescence intensity [MFI]) and additionally by counting manually under fluorescent microscope [34].

## Acknowledgements

Prof Lalita Ramakrishnan, University of Cambridge, UK, for kind donation of *Tg (mpeg1::BB)* zebrafish and all bacterial strains used in this study; and Dr Rajeeb Swain, Institute of Life Sciences, Bhubaneswar, India, for the *Tg(kdrl::EGFP)* fish Prof Paul Edelstein, University of Pennsylvania, for reviewing the manuscript.

## Author contributions

Conceptualization, SB; Methodology, SKD, RKP, SM; Validation, SKD, RKP; Formal Analysis, SB, SKD; Resources, SB, SM; Data Curation, SKD; Writing - Original Draft Preparation, SB; Writing – Review & Editing, SB, SKD, RKP, SM; Supervision, SB, SM; Project Administration, SB; Funding Acquisition, SB

